# TgATG9 is required for autophagosome biogenesis and maintenance of chronic infection in *Toxoplasma gondii*

**DOI:** 10.1101/2024.07.08.602581

**Authors:** Pariyamon Thaprawat, Zhihai Zhang, Eric C. Rentchler, Fengrong Wang, Shreya Chalasani, Christopher J. Giuliano, Sebastian Lourido, Manlio Di Cristina, Daniel J. Klionsky, Vern B. Carruthers

## Abstract

*Toxoplasma gondii* is a ubiquitous protozoan parasite that can reside long-term within hosts as intracellular tissue cysts comprised of chronic stage bradyzoites. To perturb chronic infection requires a better understanding of the cellular processes that mediate parasite persistence. Macroautophagy/autophagy is a catabolic and homeostatic pathway that is required for *T. gondii* chronic infection, although the molecular details of this process remain poorly understood. A key step in autophagy is the initial formation of the phagophore that sequesters cytoplasmic components and matures into a double-membraned autophagosome for delivery of the cargo to a cell’s digestive organelle for degradative recycling. While *T. gondii* appears to have a reduced repertoire of autophagy proteins, it possesses a putative phospholipid scramblase, TgATG9. Through structural modeling and complementation assays, we show herein that TgATG9 can partially rescue bulk autophagy in *atg9Δ* yeast. We demonstrated the importance of TgATG9 for proper autophagosome dynamics at the subcellular level using three-dimensional live cell lattice light sheet microscopy. Conditional knockdown of TgATG9 in *T. gondii* after bradyzoite differentiation resulted in markedly reduced parasite viability. Together, our findings provide insights into the molecular dynamics of autophagosome biogenesis within an early-branching eukaryote and pinpoint the indispensable role of autophagy in maintaining *T. gondii* chronic infection.

## Introduction

*Toxoplasma gondii* is an obligate intracellular protozoan parasite that infects a wide range of hosts including humans [1]. This ubiquitous pathogen is estimated to chronically infect up to one third of the global human population [2]. Chronic-stage parasites serve as reservoirs for reactivated disease under the setting of host immune dysfunction, which can lead to severe consequences such as encephalitis and blindness in affected host organs [2–4]. There have also been reports of reactivation in otherwise healthy individuals [5,6]. Current therapies against *T. gondii* fail to eliminate the chronic infection. Hence there is a critical need to better understand the mechanisms underlying *T. gondii* persistence and identify novel targets for intervention [7].

The slow-growing form of *T. gondii* (bradyzoite) that persists during chronic infection differentiates from the rapid-growing form (tachyzoite) upon external triggers such as the host immune response, restriction of nutrients, and other stresses [8]. Bradyzoites secrete proteins that form a resilient glycan-rich cyst wall that could limit nutrient access [9–11]. Nonetheless, bradyzoites can persist within cysts long term, which necessitates mechanisms in place for maintaining cellular homeostasis by degrading macromolecular material and recycling basic components. Previous studies have demonstrated that *T. gondii* parasites rely on a cysteine protease (TgCPL) residing within a plant-like vacuolar compartment (PLVAC) for degradation of proteinaceous material [12–14]. Bradyzoite mutants lacking TgCPL accumulate within the PLVAC autophagic material including organellar remnants and show reduced viability during the chronic infection [15]. This finding suggests that proteolytic turnover of cellular materials is critical for long-term parasite persistence.

Macroautophagy (hereafter, autophagy) is an important homeostatic pathway that facilitates turnover of cellular materials. The key steps involved in autophagy include initiation in response to cellular signals such as nutrient deprivation with nucleation of the phagophore, expansion of the phagophore to engulf its cargo, maturation into an autophagosome and delivery to a cell’s digestive compartment for degradative recycling [16,17]. While the molecular mechanisms of autophagy have been extensively investigated in yeast and mammalian model organisms, far less is known about the role of autophagy in early branching eukaryotes such as *T. gondii* [18,19]. Although the autophagy machinery is conserved among eukaryotes, precisely how the apicomplexan parasite *T. gondii* performs canonical autophagy with a substantially reduced repertoire of ATG (autophagy related) proteins (TgATGs) remains unclear [19]. Also, many of the discovered TgATGs including TgATG8 and its conjugation machinery play a non-canonical role in segregation of the apicoplast, a plastid organelle originating from a secondary endosymbiosis event involved in several key metabolic pathways [20–23]. However, the discovery and characterization of other TgATGs involved in canonical autophagy, specifically the early autophagy pathway, remain limited. Nonetheless, one conserved protein not involved in apicoplast segregation, TgATG9, has been reported to play an essential role in bradyzoite persistence, implying that canonical autophagy is a key survival pathway for chronic-stage parasites [24,25].

Atg9/ATG9 homologs are integral membrane proteins important for expansion of the autophagic membrane through their function as a phospholipid scramblase that moves and equilibrates phospholipids from the outer to the inner leaflet of the developing phagophore [26–30]. Atg2/ATG2 proteins transfer lipids from sites such as the endoplasmic reticulum and directly interact with Atg9/ATG9 in this step of the pathway [31–33]. Atg9/ATG9 in yeast and mammalian cells principally originate from the Golgi network and endosomal compartments with recruitment to sites of autophagosome formation upon induction of autophagy [34–36]. While these detailed characterizations of Atg9/ATG9 function and localization have been elucidated in yeast and mammalian systems, no analogous studies have been performed in *T. gondii*. Genetic ablation of TgATG9 in bradyzoites leads to impaired autophagosome production, ultrastructural abnormalities in the PLVAC, abnormal mitochondria, aberrant bradyzoite morphology *in vitro*, and decreased levels of chronic infection *in vivo* [25]. Because these phenotypes are observed with bradyzoites harboring a stable genetic knockout of *TgATG9*, it is not possible to distinguish if TgATG9 is only necessary soon after differentiation or is required to maintain the chronic infection after it is established.

In the current study, we sought direct evidence for the role of TgATG9 in autophagosome biogenesis through yeast complementation and 3D live-cell imaging. We utilized a conditional knockdown system to further determine the direct contribution of TgATG9 to autophagic flux and survival of bradyzoites after the establishment of the chronic stage.

## Results

### *TgATG9 partially rescues bulk autophagy in* atg9Δ *knockout yeast*

To determine if TgATG9 functions analogously to other Atg9/ATG9s as a putative phospholipid scramblase, we performed yeast complementation studies and assessed the rescue of bulk autophagy in *atg9Δ* knockout yeast. The TgATG9 protein contains six transmembrane domains that share homology to other Atg9/ATG9 proteins. The transmembrane domains are often termed the “core” region that confers the functionality of scrambling lipids, while the N- and C-termini have less ordered structure [26,27,30]. We performed a structural alignment of the AlphaFold2 predicted “core” of TgATG9 (aa 273-859) and *Saccharomyces cerevisiae* (Sc)Atg9 (aa 317-764), confirming that the two proteins have very similar predicted folds for their respective transmembrane regions (Figure 1A) [37]. We generated yeast expression constructs for TgATG9 driven by either the *ScATG9* promoter (*ATG9p*) or a stronger, constitutive *ZEO1* promoter (*ZEO1p*). The full-length ScAtg9 was placed under *ATG9p* expression (“Sc” in Figure 1). To account for the importance of the N- and C-termini in recruiting other autophagy proteins or in post-translational modifications, we generated a chimera (“Ch” in Figure 1) of TgATG9 and ScAtg9 by swapping the “core” of ScAtg9 with TgATG9, thus preserving the N- and C-termini as yeast sequences (Figure 1B) [38–40]. We validated the expression of all constructs by western blotting samples from unstarved and 24 h nitrogen-starved yeast (Figure 1C).

**Figure 1.**
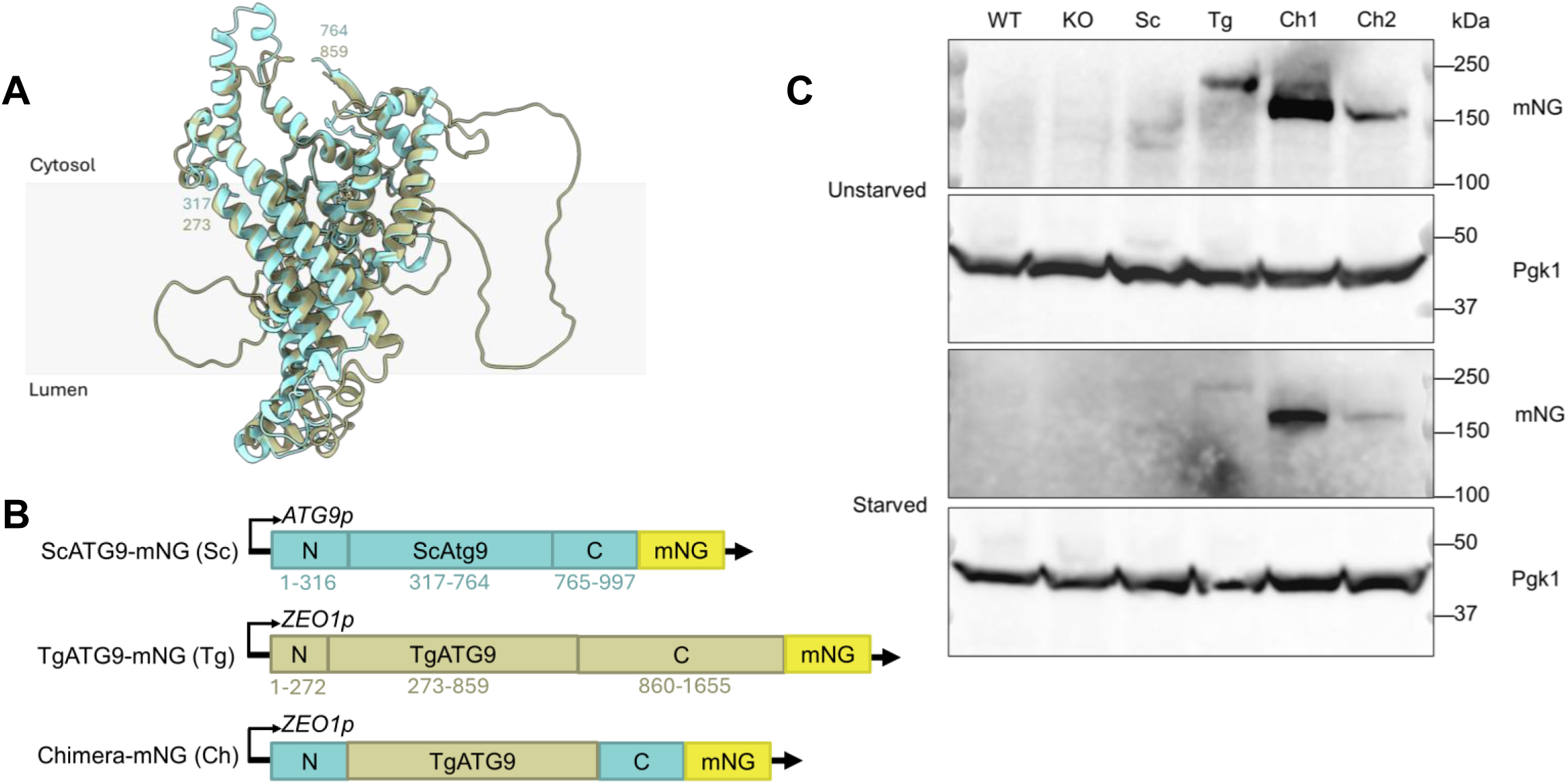
TgATG9 structural alignment with ScAtg9 and generation of constructs used for complementation assays. (**A**) Structural alignment of the AlphaFold2 predicted “core” structures of TgATG9 (olive) and ScAtg9 (cyan) using ChimeraX with a lipid bilayer depicted to represent orientation within a membrane. (**B**) Plasmids containing constructs used for yeast complementation. The yeast *ATG9* promoter (*ATG9p*) was used to drive expression of ScAtg9 while the strong, constitutive *ZEO1* promoter (*ZEO1p*) was used to drive expression of constructs containing TgATG9 sequences. All constructs had mNeonGreen (mNG) appended to the C terminus. Numbers denote amino acid position. (**C**) Western blot validation from total lysate of protein expression of constructs used for yeast complementation in unstarved and starved conditions (24 h nitrogen starvation). Pgk1 was used as a loading control. Ch1 and Ch2 are two separate isolates of the chimeric construct.

We assessed rescue of autophagic activity using the Pho8Δ60 assay and quantified the relative Pho8Δ60 activity normalized to wild-type controls [41]. Under nutrient-rich conditions (YPD medium) autophagy is not induced; however, after 4 h of nitrogen starvation (SD-N), bulk autophagy is induced and can be measured. Compared to the *atg9Δ* knockout, yeast complemented with ScAtg9 and TgATG9 showed an increase in Pho8Δ60 activity albeit not to the same levels as the wild type (Figure 2A). To further validate this finding, we assessed and quantified the level of GFP-Atg8 processing, and again showed that constructs containing *T. gondii* sequences were able to partially rescue the autophagy defect of *atg9Δ* cells (Figure 2B-C) [42]. Interestingly, there appeared to be no statistical differences between the level of rescue by TgATG9 or two independent clones of the chimera. All constructs with *T. gondii* sequences were unable to fully restore autophagy to the same levels as the wild type or yeast complemented with ScAtg9. Finally, we examined yeast viability because loss of autophagy causes a loss of viability [43,44]. To determine the effects on yeast viability, cells were plated after 6 days starvation then grown on YPD for 2 days. Wild-type cells survive starvation of this duration (Figure 2D-E). In contrast, the majority of *atg9Δ* knockout yeast plates had no colony-forming units (CFUs) observed. For the ScAtg9, TgATG9 and Ch constructs, CFUs were quantified and normalized to the wild type, which reproduced the partial rescue phenotype seen by the Pho8Δ60 and GFP-Atg8 processing assays.

**Figure 2.**
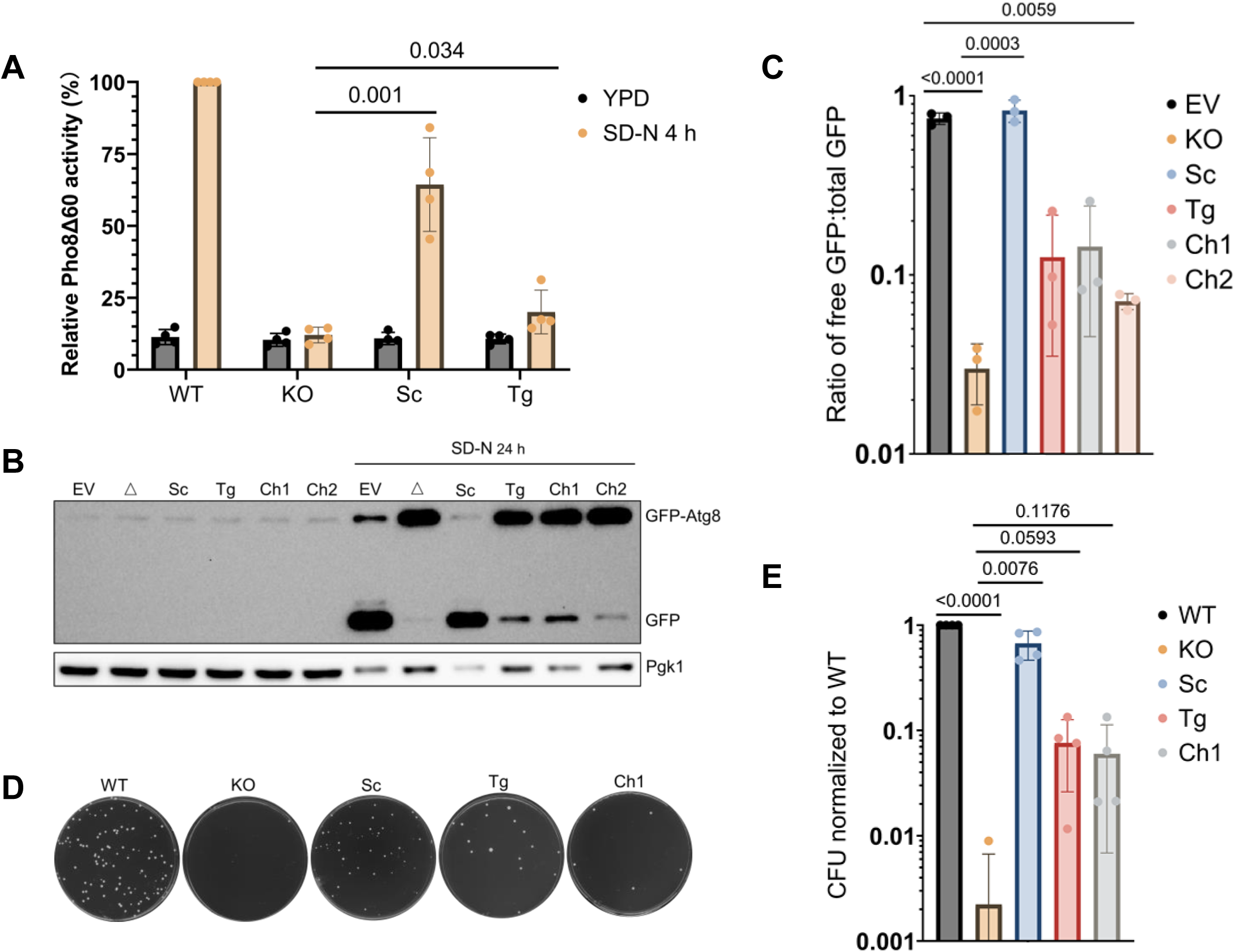
TgATG9 partially rescues bulk autophagy in *atg9Δ* knockout yeast. (**A**) Relative Pho8Δ60 activity normalized to wildtype (WT) controls for yeast *atg9Δ* (KO) and KO complemented with ScAtg9 (Sc) or TgATG9 (Tg) in nutrient-rich (YPD) or nitrogen-starvation (SD-N 4 h) conditions (n=4 biological replicates). Bars represent mean ± SD. Statistical analysis was done using a Student’s *t*-test with p-values denoted above bars. (**B, C**) GFP-Atg8 processing assay. Representative western blot (B) depicting GFP-Atg8 or free GFP in unstarved or starved conditions (SD-N 24 h). WT control with empty vector (EV) or KO (Δ) were complemented with plasmids containing ScAtg9 (Sc), TgATG9 (Tg), or chimera (Ch1 and Ch2) constructs. Pgk1 was used as a loading control. (C) Ratio of free GFP:total GFP quantified from (B). Bars represent mean ± SD. Statistical analysis was done using a one-way ANOVA followed by a Student’s *t*-test with p-values denoted above bars (n=3 biological replicates). (**D, E**) CFU viability assay. Representative images from 100 μL (1 OD diluted to 10^-4^) plated cells after 6 days of nitrogen starvation then grown in YPD for 2 days. (E) Quantification of (D) normalized to WT controls. Bars represent mean ± SD. Statistical analysis was done using a one-way ANOVA followed by a Student’s *t*-test with p-values denoted above bars (n = 3 biological replicates).

To confirm that TgATG9 is being trafficked to the phagophore assembly site/PAS, we transformed ScAtg9 and TgATG9 into *atg1Δ atg9Δ* RFP-Atg8-expressing yeast. Atg1 is required for retrograde transport of Atg9; in *atg1Δ* strains, Atg9 tends to accumulate at the phagophore assembly site [45]. We found that within 1 h of nitrogen starvation both proteins colocalized with RFP-Atg8 in the *atg1Δ* background, suggesting proper localization (Figure S1) [46]. However, qualitatively, there appeared to be more clustered RFP-Atg8 puncta present when ScAtg9 was introduced compared to TgATG9, especially after 3 h of starvation, which is congruent with the partial phenotypes we have observed. These findings could indicate defects in proper cycling of TgATG9, but interrogating this further is beyond the scope of this study. Additionally, while the main scramblase activity of Atg9 is conferred by its transmembrane region, the N- and C-termini contain phosphorylation sites necessary for Atg9 activity or stability [40,47]. We searched the *Toxoplasma* database (ToxoDB) and found that TgATG9 contains six phosphorylation sites based on published phosphoproteomics studies (Figure S2A-B) [48,49]. We generated tdTomato-TgATG8-expressing TgATG9 phosphorylation mutant parasites (S165A, S933A, S1316A, S1332A, S1377A, S1482A) and assessed autophagy function by using a fluorescent dye (CytoID) that stains autolysosomes. CytoID staining was performed after treatment with the cathepsin protease L (TgCPL) inhibitor LHVS to allow for autophagic material to accumulate for visualization [15,50]. Both wild-type (ME49) and TgATG9 phosphorylation mutant parasites accumulated CytoID-positive puncta after LHVS treatment, suggesting that TgATG9 phosphorylation status was not critical for autophagic function (Figure S2C).

Taken together, our data demonstrate that while TgATG9 is dissimilar in many ways to ScAtg9 including size and post-translational modifications, it contains a core transmembrane structure that likely confers phospholipid scramblase function that can partially rescue bulk autophagy in *atg9Δ* knockout yeast.

### Conditional knockdown of TgATG9 disrupts bradyzoite autophagy

We previously demonstrated the importance of TgATG9 in *T. gondii* autophagy using a parasite line harboring a stable *TgATG9* genetic knockout. However, we could not rule out indirect effects or delayed effects from the complete absence of TgATG9 on bradyzoite biology [25]. To provide direct evidence that TgATG9 is critical and specific for autophagy, we generated a conditional knockdown mutant of TgATG9 using the auxin-inducible degron (AID) system that confers post-translational reversible regulation [51]. In parasites expressing the heterologous F-box protein TIR1, TgATG9 was endogenously tagged at the C terminus with a minimal auxin-inducible degron (mAID) fused to the mNeonGreen fluorophore (Figure 3A) [52]. Upon addition of auxin (indole-3-acetic acid, IAA), we observed efficient conditional knockdown of TgATG9-mAID in *in vitro* bradyzoites after 24 h by live fluorescence microscopy and western blotting (Figure 3B-C). Next, we assessed the impact of conditional knockdown of TgATG9-mAID on bradyzoite autophagy by differentiating *in vitro* bradyzoites for 7 days followed by treatment with IAA or vehicle control with or without LHVS for 3 days prior to CytoID staining. Bradyzoites of the TIR1 parental strain accumulated CytoID-positive autolysosomes in the presence or absence of IAA treatment, as expected (Figure 3D). Similarly, without IAA treatment TgATG9-mAID parasites accumulated CytoID-positive autolysosomes. However, TgATG9-mAID bradyzoites treated with IAA showed markedly reduced CytoID-positive staining of autolysosomes (Figure 3D). These findings provide direct evidence that TgATG9 is important for bradyzoite autophagy and establish a conditional knockdown system to allow us to assess the direct impacts of acute disruptions to this pathway.

**Figure 3.**
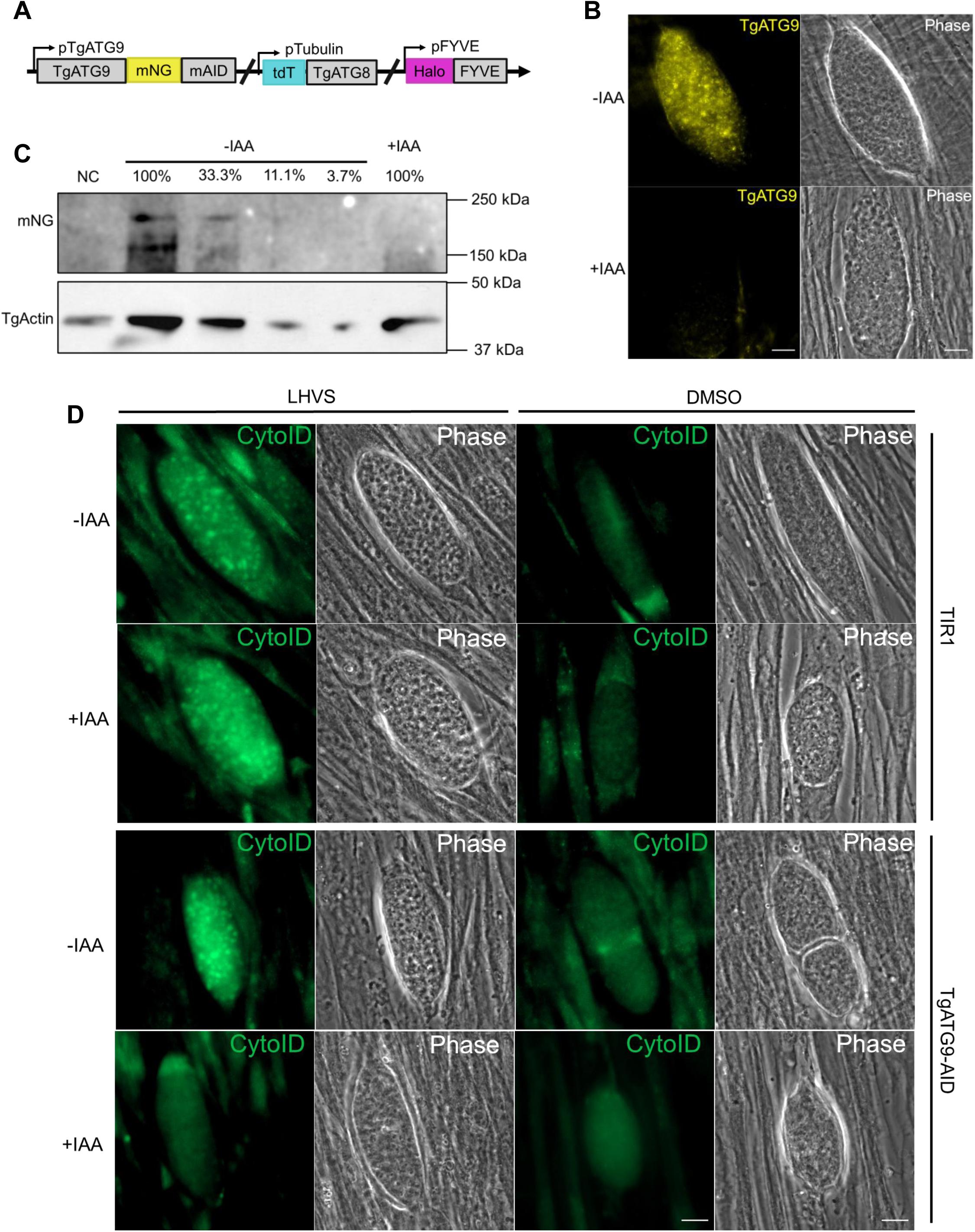
Conditional knockdown (KD) of TgATG9 disrupts bradyzoite autophagy. (**A**) Endogenous tagging of TgATG9 with mNeonGreen (mNG) and the minimal auxin inducible degron (mAID) was achieved with CRISPR-Cas9 and homology-directed repair. tdTomato (tdT)-TgATG8 was subsequently introduced as an exogenous copy by insertion into the tubulin locus. Endogenous tagging of TgFYVE with HaloTag (Halo) was subsequently achieved with CRISPR-Cas9 and homology-directed repair to generate triple-fluorescent parasites. (**B**) Fluorescent-imaging of *in vitro* TgATG9-mAID differentiated cysts treated with IAA or vehicle control for 24h. Scale bars: 10 μm. (**C**) Western blot from total bradyzoite lysates of conditional KD of TgATG9-mAID with IAA or vehicle control for 24 h. Serial dilution of vehicle treated parasites were used for relative quantification of protein levels. TgActin was used as a loading control. (**D**) Autolysosome staining (CytoID) of *in vitro* TIR1 or TgATG9-mAID differentiated cysts treated with or without LHVS and IAA or vehicle control for 72 h. Scale bars: 10 μm. (n = 3 biological replicates, representative images shown).

### TgATG9 is required for autophagosome biogenesis and proper autophagic flux

To define the role of TgATG9 in autophagic flux, we first confirmed the subcellular localization of TgATG9 within bradyzoites by performing immunofluorescence staining and quantified colocalization to known autophagy markers in fixed samples. TgATG9 colocalized with TgCPL (PLVAC marker) and TgATG8 (Figure S3). We included staining for ATRX1, an apicoplast marker, as a control and showed that TgATG9 colocalizes more significantly to autophagic structures rather than the apicoplast, in agreement with previous studies performed in tachyzoites [24]. While these data provide insight into the location of TgATG9 within bradyzoites, we recognize that they are only two-dimensional snapshots of an intrinsically dynamic process. Therefore, we wanted to assess the dynamics of this pathway by live microscopy.

We hypothesized that the acute disruption of bradyzoite autophagy would result in altered dynamics between TgATG9, TgATG8, and the PLVAC. Using our TgATG9-mAID strain, which also has mNeonGreen fused to TgATG9, we further generated a triple-fluorescent parasite strain by ectopically expressing tdTomato-TgATG8 as a marker of autophagosomes and endogenously HaloTag-tagging TgFYVE as a marker for the PLVAC (Figure 3A) [53,54]. Given that there are many bradyzoites tightly packed and randomly oriented within an intracellular cyst, we aimed to capture autophagosome dynamics in a 3-dimensional manner over a time course of several minutes utilizing lattice light sheet microscopy (LLSM) to image live, *in vitro* differentiated cysts. LLSM permits rapid imaging of several planes over a given time course and therefore is well suited to capture cellular dynamics within the 3-dimensional structure of cysts with minimal photobleaching [55–57].

Using our triple-fluorescent parasites, we assessed the direct effects of conditional knockdown of TgATG9-mAID by LLSM (Figure 4A). Parasites were differentiated into bradyzoites *in vitro* for 7 days prior to treatment with IAA or vehicle control for 24 h. We then imaged intracellular cysts containing bradyzoites over the course of 5 min. First, we observed a reduction in TgATG9 puncta in the IAA-treated cysts, validating the effectiveness of our conditional knockdown (Figure 4B, Video S1). Within untreated cysts, we observed examples of the spatiotemporal dynamic associations between TgATG9, TgATG8, and the PLVAC resembling autophagosome trafficking to the digestive organelle or autophagosome fusion with the PLVAC (Figure 4C, Video S2). We quantified the degree of colocalization between the various markers, accounting for the knockdown of TgATG9, reported as a ratio of co-occurrence. Globally, on a per cyst level, IAA-treated bradyzoites displayed a significant reduction in the co-occurrence of TgATG9 or TgATG8 with the PLVAC, suggesting decreased autophagic flux due to impaired delivery of autophagosomes to the PLVAC compared to vehicle-treated controls (Figure 4D); there was essentially no difference in the co-occurrence between TgATG9 and TgATG8. Using Arivis Vision4D software, we generated volumetric renderings of the three markers and calculated their mean object volumes within a given cyst. IAA-treated parasites had significantly smaller TgATG9 and TgATG8 puncta compared to vehicle-treated controls, indicating an impairment in the biogenesis and enlargement of autophagosomes (Figure 4E). Additionally, TgATG9 knockdown bradyzoites had significantly smaller PLVACs, which is consistent with decreased delivery of autophagic material to the PLVAC.

**Figure 4.**
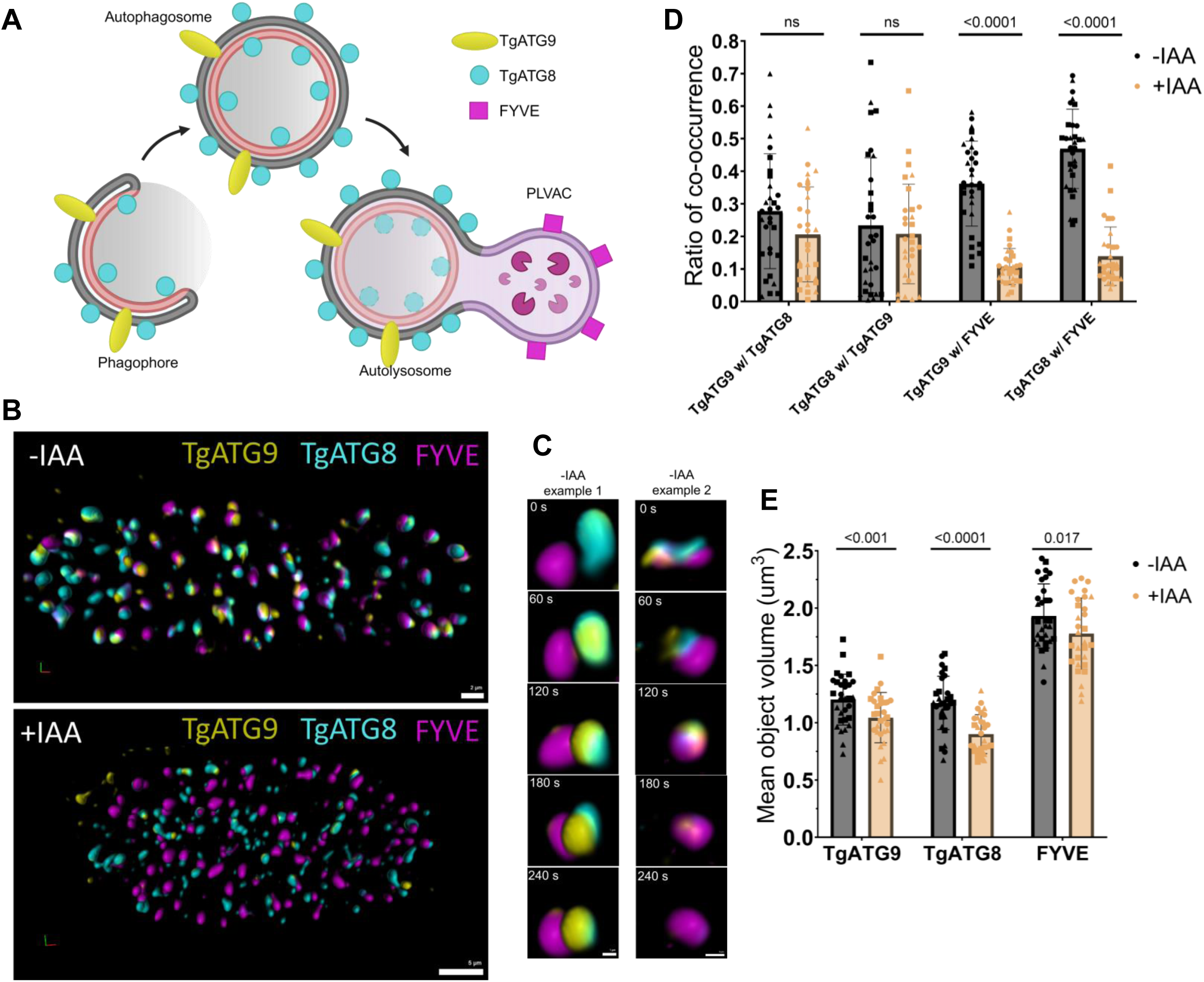
Conditional KD of TgATG9 disrupts dynamic associations between autophagosomes and the PLVAC in bradyzoites. (**A**) Schematic of fluorescent markers monitored in lattice light sheet microscopy (LLSM) experiments created with BioRender. (**B**) Representative *in vitro* differentiated cysts imaged with LLSM treated with IAA or vehicle control for 24 h. Scale bars: 2 μm (top) and 5 μm (bottom). (**C**) Representative examples of dynamic associations between TgATG9, TgATG8, and FYVE imaged with LLSM in untreated parasites (-IAA). Scale bars: 1 μm (left) and 5 μm (right). Volumetric renderings for (B, C) generated with Arivis Vision4D software. (**D, E**) Quantification of the ratio of co-occurrence and mean object volumes among TgATG9, TgATG8, and TgFYVE from LLSM. Bars represent mean ± SD. Statistical analysis was done using paired Student’s *t*-test with p-values denoted above bars. (n=3 biological replicates, ≥10 cysts’ total volumes were imaged per treatment condition over 5 min per replicate).

Thus, our findings support the direct role of TgATG9 in autophagosome biogenesis in bradyzoites whereby acute disruption of the autophagy pathway results in decreased autophagic flux through altered dynamics between autophagosomes and delivery of autophagic material to the PLVAC.

### Bradyzoite autophagy is required for maintenance of chronic infection in vitro

While we have demonstrated that bradyzoites with a stable genetic knockout of *TgATG9* have reduced viability, we cannot rule out any accumulated effects from the absence of TgATG9 in establishment or persistence of chronic infection [25]. Although TgATG9 is not essential for normal tachyzoite growth, TgATG9 knockout tachyzoites have reduced viability after extracellular stress and have decreased virulence during mouse infection [24]. To pinpoint the role of TgATG9 and autophagy in parasite persistence specifically within the bradyzoite stage, we assessed bradyzoite viability after conditional knockdown of TgATG9 post-differentiation *in vitro*. TIR1 or TgATG9-mAID parasites were differentiated into bradyzoites for 7 days then treated with constant IAA or vehicle control for an additional 7 days with daily media changes. After the 7 days of treatments, bradyzoites were harvested from cultured cysts using pepsin and their viability was assessed by plaque assay with qPCR normalization of parasite genomes (Figure 5A) [58]. IAA treatment had no effect on the viability of TIR1 bradyzoites, while IAA-treated TgATG9-mAID parasites had significantly reduced viability (Figure 5B). Our results emphasize the importance of TgATG9 and autophagy in sustaining bradyzoite survival and clarify their critical role in the maintenance of chronic infection.

**Figure 5.**
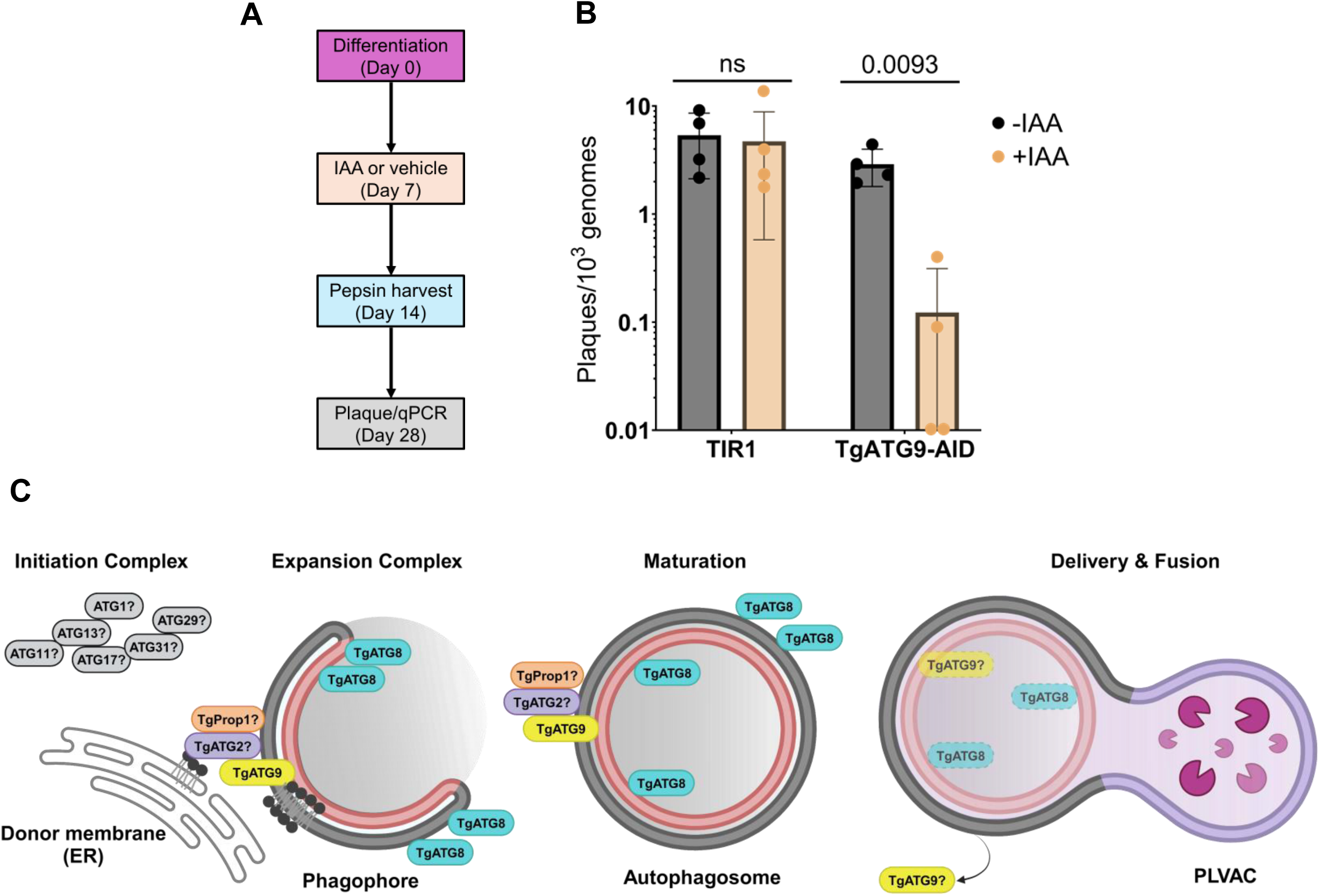
*T. gondii* bradyzoite autophagy is required for maintenance of chronic infection *in vitro*. (**A**) Schematic of bradyzoite viability assay. Parasites were differentiated in alkaline-stress media for 7 days prior to initiation of treatment with IAA or vehicle control for another 7 days with daily media changes. Bradyzoites were harvested after 7 days treatment (Day 14) via pepsin harvest and directly used for plaque assay and genomic DNA extraction for qPCR. Plaques were quantified after 14 days (Day 28). (**B**) Viability of *in vitro* 7-days differentiated bradyzoites after 7 days treatment with IAA or vehicle control reported as number of plaques normalized to number of parasite genomes. Bars represent mean ± SD. Statistical analysis was done using paired Student’s *t*-test with p-values denoted above bars. (n=4 biological replicates). (**C**) Model of the role of TgATG9 in autophagosome biogenesis in *T. gondii* canonical autophagy. Autophagy is initiated through complex signals including response to cellular stress or nutrient deprivation that thereby activates an initiation complex comprised of key proteins yet to be discovered and characterized in this system. Downstream of the initiation complex, TgATG9 is the only known transmembrane autophagy protein that resides on the developing phagophore that likely nucleates a putative membrane expansion complex with potential interactions with other key proteins involved in this process (putative TgATG2 and TgATG18/TgProp1 have yet to be discovered or characterized in this context). Lipids from a donor membrane such as the ER are transferred onto the phagophore for membrane expansion whereby TgATG9 equilibrates donated lipids on the phagophore membrane. TgATG8 also gets lipidated onto the autophagosome membrane during maturation. The mature autophagosome gets delivered to the parasite’s digestive organelle (PLVAC) and fuses with the PLVAC for delivery of autophagic cargo that will undergo degradative and recycling processes within the PLVAC lumen. It remains unknown whether TgATG9 gets degraded within the PLVAC or whether *T. gondii* autophagy possesses a mechanism for the cycling of TgATG9 like other model systems. Schematic created with BioRender.

## Discussion

While it has been established that autophagy is critical for *T. gondii* bradyzoites, the early steps of the pathway remain poorly characterized. Here, we expand our understanding of an important component of this pathway, TgATG9, by providing additional evidence for its role in autophagosome biogenesis. We suggest that TgATG9 contributes to the expansion of the phagophore by equilibrating phospholipids across the lipid bilayer (Figure 5C). Although it is a much larger protein than ScAtg9 and has limited homology apart from the core domain, TgATG9 was able to partially rescue bulk autophagy in *atg9Δ* knockout yeast, which we validated across three independent complementation assays. We investigated the spatiotemporal dynamics of TgATG9 in bradyzoite autophagy through LLSM, finding that TgATG9 is key to proper autophagosome formation and delivery to the PLVAC. As a putative phospholipid scramblase, TgATG9 likely requires collaboration with other TgATGs that participate in the membrane expansion complex such as a lipid transfer protein (a hypothetical TgATG2) and phosphatidylinositol-3-phosphate/PtdIns3P-binding protein (TgProp1, an Atg18 ortholog). Whether *T. gondii* forms such a membrane expansion complex will require the discovery and further characterization of candidate proteins. The fate of TgATG9 upon autophagosome fusion with the PLVAC also remains elusive. TgATG9 may be degraded within the PLVAC or potentially cycled through unknown mechanisms. Additionally, the absence of known initiators or regulators of *T. gondii* autophagy further contributes to our incomplete understanding of this pathway.

Our yeast complementation studies demonstrated that expression of either full-length TgATG9 or a chimera of TgATG9/ScAtg9 resulted in partial rescue of bulk autophagy. There are several possible explanations for the lack of full rescue. We determined the boundaries for the “core” region of TgATG9 and ScAtg9 based on a protein-protein BLAST search which may not have accurately mapped the true “core” region of transmembrane domains and important flanking residues [59]. Atg9/ATG9 proteins are known to form homotrimers with each protomer’s transmembrane domains contributing to formation of a central cavity for scrambling lipids, a function that can be disrupted if interactions within the homotrimeric interface are altered [26,27,30,60]. In yeast, Atg9 is post-translationally modified through phosphorylation in Atg1-dependent and -independent manners. Such modifications regulate the cycling of Atg9 to phagophore assembly sites or protein levels of Atg9, thus controlling rates of autophagosome formation [40,47,61]. We considered the importance of these post-translational modifications when designing the chimera, thereby preserving phosphorylated residues. However, we found that preservation of these sites did not improve the partial rescue phenotype observed compared to yeast complemented with the full-length TgATG9. We tested the importance of TgATG9 phosphorylation in *T. gondii* and found that the annotated phosphorylation sites are not important in bradyzoite autophagy. However, these sites are based on phosphoproteomic data from tachyzoites; hence, additional studies are needed to determine the extent of TgATG9 phosphorylation specifically in bradyzoites [49]. Nonetheless, our assessment of TgATG9 phosphorylation provides some insights into potential key differences in how autophagy may be regulated in *T. gondii.* This is particularly intriguing because an initiating kinase such as Atg1 or ULK1 has not been discovered in *T. gondii* [18,19]. Although our studies primarily focused on regions of TgATG9 and ScAtg9 with shared homology, consideration of dissimilar regions may uncover novel features which could expand our understanding of the evolution of autophagy among eukaryotes.

We localized TgATG9 to autophagic structures marked with TgATG8 (autophagosomes) and TgFYVE (PLVAC) within bradyzoites and utilized LLSM to visualize these dynamic associations that became disrupted upon conditional knockdown of TgATG9. The decreased size of autophagosomes (TgATG9 and TgATG8) along with decreased colocalization between autophagosomes and the PLVAC indicate that TgATG9 is crucial for autophagosome biogenesis. Under normal conditions, we observed examples of autophagosome delivery or fusion with the PLVAC occurring within a few minutes, consistent with recently reported spatiotemporal observations of autophagosome formation [62]. However, our LLSM studies lacked single-cell resolution and to definitively determine or quantify the autophagosome dynamics within a single bradyzoite would require additional labeling of markers at the parasite membrane or inner membrane complex. Despite this, we provide the first spatiotemporal characterization of autophagosome biogenesis in *T. gondii* and demonstrate the feasibility of monitoring live subcellular dynamics in bradyzoites.

We directly determined the role of autophagy in maintenance of chronic infection and showed that conditional knockdown of TgATG9 post-differentiation results in reduced bradyzoite viability. These findings strengthen our conclusion that autophagy is specifically important for bradyzoite survival after chronic infection is already established. This study therefore addresses any potential confounders of altered bradyzoite biology that were observed with the stable *TgATG9* genetic knockout including altered mitochondrial morphology, a phenotype observed as early as 6 days into *in vitro* differentiation. *TgATG9* knockout bradyzoites also display impaired replication and division along with a disorganized inner membrane complex [25]. While we did not directly investigate these changes with our conditional knockdown of TgATG9, we can infer that such changes did not come into play during the establishment of chronic infection in our experiments. Future studies could extend our *in vitro* findings into an *in vivo* mouse model to provide proof of concept that bradyzoite autophagy is selectively targetable to potentially curb chronic infection. Furthermore, the mechanism by which bradyzoites are dying remains to be elucidated. Additional studies are required to determine how disrupting autophagy affects bradyzoite biology, perhaps due to accumulation of damaged organelles, altered metabolism, or activated cellular death pathways. Our TgATG9 conditional knockdown mutant can be used in future metabolomic or proteomic studies for this purpose. While several other pathways have been reported to be critical to bradyzoite survival including storage and utilization of amylopectin granules, maintaining structural integrity of the cyst wall, and apicoplast-related functions, our results underscore the importance of autophagy in this stage of *T. gondii* [8,63–66].

Taken together, our study advanced understanding of the direct contributions from TgATG9 to autophagosome biogenesis in bradyzoites and maintenance of chronic infection. Further characterization of the autophagy pathway, particularly the poorly understood early steps, will provide valuable insights into the evolution, mechanism, and regulation of this indispensable process that can be strategically manipulated to target chronic infection.

## Materials and Methods

### Structural prediction and alignment of TgATG9 and ScAtg9

A predicted structure of the TgATG9 (TGME49_260640) monomer was generated using AlphaFold2 implemented in ColabFold v1.5.5 on the Google Colab platform (https://colab.research.google.com/github/sokrypton/ColabFold/blob/main/AlphaFold2.ipynb) [37,67]. The full-length sequence of TgATG9 (1-1655) was used as the search input. Five predicted models were generated and ranked based on pLDDT and PAE scores. The “core” region from the top-ranked model of TgATG9 (273-859) was structurally aligned with the “core” region of a monomer of the AlphaFold predicted model of *Saccharomyces cerevisiae* Atg9 (317-764) (https://alphafold.ebi.ac.uk/entry/Q12142) using Matchmaker structure analysis in UCSF ChimeraX version 1.7.1 [68]. A figure of the aligned predicted structures was prepared in ChimeraX.

### Yeast strains, media, and growth conditions

All yeast *Saccharomyces cerevisiae* strains used in this study are listed in Table S1 [47,69–71]. Gene deletions and chromosomal tagging were performed using standard methods [72]. Yeast cells were cultured at 30°C in rich medium (YPD; 1% yeast extract [Formedium, YEA04], 2% peptone and 2% glucose) as nutrient-rich conditions. To induce autophagy, cells in mid-log phase (OD600 = 0.4-0.8) were shifted from YPD to nitrogen starvation medium (SD-N; 0.17% yeast nitrogen base without ammonium sulfate or amino acids [Formedium, CYN0501], containing 2% glucose) for the indicated times.

### Generation of plasmids for yeast complementation

To generate the plasmids for the yeast autophagy complementation experiments, plasmid pRS406-ATG9p-ScAtg9-mNeonGreen was constructed by amplifying the *ATG9* promoter region (−1000-0) and *ATG9* ORF by PCR from genomic DNA. The PCR product was inserted into the pRS406-mNeonGreen plasmid using Gibson Assembly. Of note, the *ZEO1* promoter region (−550-0) was amplified by PCR from genomic DNA and then the PCR product was used to replace the *ATG9* promoter region on the pRS406-ATG9p-ScAtg9-mNeonGreen plasmid by Gibson Assembly to construct the pRS406-ZEO1p-ScAtg9-mNeonGreen plasmid to facilitate other plasmid construction. Plasmid pRS406-ZEO1p-TgAtg9-mNeonGreen plasmid was constructed by replacing the *ATG9* ORF region on the pRS406-ZEO1p-ScAtg9-mNeonGreen plasmid with the *TgATG9* ORF region amplified from genomic DNA. Similarly, plasmid pRS406-ZEO1p-Sc/TgAtg9chimera-mNeonGreen plasmid was constructed by replacing the “core” *ATG9* ORF of ScAtg9 region on the pRS406-ZEO1p-ScAtg9-mNeonGreen plasmid with the “core” *TgATG9* ORF region amplified from genomic DNA. Before transformation, pRS406-based plasmids were cut with BstBI at 65°C.

### Yeast complementation assays

#### Pho8Δ60 assay

Pho8Δ60 assays were performed as previously described [41].

#### GFP-Atg8 processing assay

GFP-Atg8 processing assays were performed as previously described [73]. Rabbit anti-Pgk1 antiserum is a generous gift from Dr. Jeremy Thorner (University of California, Berkeley; 1:1000). Other antibodies were from the following sources: mouse anti-YFP, which detects GFP in our study (Clontech, 63281; 1:5000), goat anti-rabbit IgG secondary antibody (Fisher, ICN55676; 1:10000) and rabbit anti-mouse IgG secondary antibody (Jackson, 315-035-003; 1:10000). The blot was imaged using ChemiDoc Touch imaging system (Bio-Rad, 1708370) and quantified using Bio-Rad Image Lab software.

#### CFU viability assay

Yeast cells were grown in YPD to mid-log phase and then shifted to SD-N medium for nitrogen starvation. After 6 days starvation, 1 OD600 units of cells were collected, resuspended in 1 mL of SD-N medium (1 OD600/mL) and then subjected to serial dilution. 100 μL of 10^-4^ diluted samples were spread onto YPD plates and incubated at 30°C for 2 days before imaging.

### Parasite and host cell culture

All *T. gondii* parasites were cultured in human foreskin fibroblasts (HFFs, Hs27) obtained from the American Type Culture Collection (ATCC-CRL-1634). Tachyzoite growth was maintained at 37°C under 5% CO_2_ in D10 medium, comprising DMEM (Fisher Scientific, 10-013 cv), 10% heat-inactivated bovine calf serum (Cytiva, SH30087.03), 2 mM L-glutamine (Corning, 25-005-Cl), and 50 U/mL of penicillin-streptomycin (Gibco, 15070063). For all bradyzoite differentiation, tachyzoites were mechanically lysed from infected HFF monolayers via scraping and syringing (20- and 25-gauge needles) and allowed to invade fresh monolayers of HFFs for 24 h at 37°C under 5% CO_2_ in D10 medium. The following day, D10 medium was replaced with alkaline-stress medium consisting of RPMI-1640 without NaHCO_3_ (Cytiva, SH30011.02) supplemented with 3% heat-inactivated fetal bovine serum (Cytiva, SH30396.03), 50 mM HEPES (Sigma-Aldrich, H3375), 50 U/mL of penicillin-streptomycin and adjusted to pH 8.2-8.3 with NaOH. All bradyzoites were cultured at 37°C under ambient CO_2_ conditions with daily medium changes to ensure alkaline pH [74].

### T. gondii *transfection*

Intracellular tachyzoites were mechanically lysed from infected HFF monolayers via scraping and syringing (20- and 25-gauge needles). Parasites were pelleted and resuspended in cytomix transfection buffer (2 mM EDTA, 120 mM KCl, 0.15 mM CaCl_2_, 10 mM K_2_HPO_4_/KH_2_PO_4_, 25 mM HEPES, 5 mM MgCl_2_, pH 7.6). For each transfection, guide RNA plasmids, homology-direct repair templates, or linearized plasmids were precipitated with ethanol and resuspended in cytomix transfection buffer. The DNA mixture was combined with pelleted parasites followed by electroporation in 0.4-cm cuvettes (Bio-Rad, 1652081), utilizing GenePulser Xcell with PC and CE modules (Bio-Rad, 1652660), and configured with the following parameters: 2400 V voltage, 25 μF capacitance, 50 Ω resistance.

### T. gondii *strain generation*

Primers and oligos were synthesized either by IDT or Sigma-Aldrich. All guide RNAs were generated by substituting the original 20 base pair guide RNA sequence on the plasmid pCas9/sgRNA/Bleo [75] with desired 20 base pair guide RNA sequence, using the Q5® Site-Directed Mutagenesis Kit (New England Biolabs, E0554S). Homology-directed repair templates were PCR amplified using the CloneAmp™ HiFi PCR Premix (Takara Bio, 639298). Guide RNA and repair template sequences are listed in Table S1.

### *Generation of the TgATG9-mAID* T. gondii *strain*

Staring with the TIR1-expressing ME49 *T. gondii* strain (ME49/TIR1) [52], TgATG9 was endogenously tagged at the C terminus to generate ME49/TIR1/TgATG9-mAID (TgATG9-mAID) parasites. A guide RNA targeting the C terminus of TgATG9 near the stop codon was generated using oligos P1/P2. Homology-directed repair template was generated using oligos P3/P4 for PCR amplification of pGL015 [76] containing the Xten linker, V5 epitope (Invitrogen, 37-7500), mNeonGreen, minimal AID, and Ty epitope. ME49/TIR1 parasites were co-transfected with 100 µg guide RNA and 50 µg repair template. At 48 h after transfection, positively transfected parasites were selected using 5 µg/mL phleomycin (Invitrogen, NC9198593) prior to isolating clones by limiting dilution. Individual parasite clones were validated by PCR amplification to confirm presence of the mAID tag using P5/P6 and live imaging for mNeonGreen fluorescence.

### *Generation of the triple-fluorescent* T. gondii *strain used in LLSM*

Starting with the ME49/TIR1/TgATG9-mAID (TgATG9-mAID) *T. gondii* strain, a triple-fluorescent strain was generated by ectopic expression of tdTomato-TgATG8 and endogenous tagging of HaloTag-TgFYVE at the N-terminus. Plasmids containing pTubulin-tdTomato-TgATG8-BLE [15] were linearized with PmeI and 100 µg was transfected into TgATG9-mAID parasites for single crossover integration into the tubulin locus. tdTomato^+^ parasites were enriched by sorting 48 h post-transfection prior to isolating clones by limiting dilution and phleomycin selection. Individual parasite clones (ME49/TIR1/TgATG9-mAID/tdTomato-TgATG8) were validated by live imaging for tdTomato fluorescence. In parallel, the parental ME49/TIR1 *T. gondii* strain was also co-transfected with the same amount of linearized plasmids to generate ME49/TIR1/tdTomato-TgATG8 parasites to be used as controls in LLSM.

Subsequently, a homology-directed repair template was generated using oligos P7/P8 for PCR amplification of a FLAG-HaloTag-containing plasmid (phage-ubc-flag-halo-baCDS-baUTR1-24xMS2V5-wpre, Louis M. Weiss lab [Albert Einstein College of Medicine] unpublished). Double-fluorescent (ME49/TIR1/TgATG9-mAID/tdTomato-TgATG8) parasites were co-transfected with 100 µg of a guide RNA targeting the N terminus of TgFYVE (TGME49_237870) [54] and 50 µg repair template. At 48 h after transfection, positively transfected parasites were selected using 5 µg/mL phleomycin prior to isolating clones by limiting dilution. Individual parasite clones (ME49/TIR1/TgATG9-mAID/tdTomato-TgATG8/HaloTag-FYVE) were validated by PCR amplification to confirm presence of the HaloTag using P9/P10. In parallel, the ME49/TIR1/tdTomato-TgATG8 *T. gondii* strain was also co-transfected with the same amounts of guide RNA and repair template to generate ME49/TIR1/tdTomato-TgATG8/HaloTag-FYVE parasites to be used as controls in LLSM.

### Widefield fluorescence microscopy

For live microscopy used to validate conditional knockdown of TgATG9-mAID, parasites were differentiated into bradyzoites for 7 days in 35-mm dish, No. 1.5 coverslip, 14-mm diameter, tissue culture dishes (MatTek, P35G-1.5-14-C) in phenol red free alkaline-stress medium consisting of modified RPMI-1640 without phenol red or NaHCO_3_ (Sigma-Aldrich, R8755) supplemented with 3% heat-inactivated fetal bovine serum, 50 mM HEPES, 50 U/mL of penicillin-streptomycin and adjusted to pH 8.2-8.3 with NaOH. After 7 days differentiation, bradyzoites were treated with 500 µM IAA (Sigma-Aldrich, I2886) or an equal volume ethanol vehicle control for 24 h for conditional knockdown of TgATG9-mAID. Widefield microscopy images were taken on a Zeiss Axio Observer Z1 inverted microscope at 40× and analyzed using Zen 3.7 blue edition software. For imaging of fixed samples for all CytoID experiments, images were taken at 63× and analyzed using Zen 3.7 blue edition software.

### Immunoblotting

For validation of TgATG9 knockdown by immunoblot, TgATG9-mAID parasites were differentiated into bradyzoites for 7 days grown in alkaline-stress medium (ambient CO_2_) with daily media changes. On day 7, bradyzoites were treated with 500 µM IAA (Sigma-Aldrich, I2886) or ethanol vehicle control for 24 h. Bradyzoites were harvested via scraping, syringing (20- and 25-gauge needles), pepsin treatment (0.026% pepsin [Sigma-Aldrich, P7000] in 170 mM NaCl and 60 mM HCl, final concentration), and filtration. Bradyzoites were enumerated and lysed with RIPA buffer (Thermo Scientific, 89900) supplemented with cOmplete Mini Protease Inhibitors cocktail (Roche, 11836153001) for 15 min on ice with gentle rocking. Lysates were centrifuged at 4°C for 10 min at 20,000 x g. Lysates were supplemented with 5X SDS-PAGE sample buffer and 10% β-mercaptoethanol, resulting in a final concentration of 1X SDS-PAGE buffer and 2% β-mercaptoethanol. The final concentration was ∼4×10^7^ bradyzoites per 25 µL and designated as 100% for loading.

Lysates were subjected to SDS-PAGE using a gradient 4-12% NuPAGE™ Bis-Tris gel (Invitrogen, NP0321) and transferred onto 0.45-µm nitrocellulose membranes (Bio-Rad, 1620115) with Trans-Blot® SD semi-dry transfer cell (Bio-Rad, 1703940) for 45 min at 18 V at room temperature. Following transfer, membranes were blocked with 5% milk in phosphate-buffered saline (PBS [Gibco, 21600010])-T (PBS with 0.05% Triton X-114 [Sigma-Aldrich, X114] and 0.05% Tween-20 [Fisher Scientific, 170-6531]) for 30 min at room temperature. Primary antibodies were diluted in 1% milk in PBS-T and applied to membranes overnight at 4°C. Primary antibodies used include mouse anti-mNeonGreen (Chromotek, 32F6; 1:1000), rabbit anti-TgActin (Sibley lab, Washington University in St. Louis; 1:20,000). After primary antibody incubation, membranes were washed 3 times with PBS-T before incubation with HRP-conjugated secondary antibodies (Jackson ImmunoResearch Laboratories, 115-035-146; 1:5000) for 1 h at room temperature. Proteins were detected using SuperSignal™ West Pico PLUS Chemiluminescent Substrate or Femto Maximum Sensitivity Substrate (Thermo Fisher, 1863096 or 34095). The Syngene PXi6 imaging system with Genesys (v1.8.2.0) software was used to detect signals, and Fiji (v2.9.0/1.53t) [77] was used for quantification.

### CytoID staining of autolysosomes

*T. gondii* tachyzoites were differentiated into bradyzoites in 6-well tissue culture plates with 22 mm x 22 mm No. 1.5 coverslips (Globe Scientific, 1404-15) for 7 days then treated with 1 µM LHVS (Sigma-Aldrich, SML2857) or equal volume DMSO for 3 days. For knockdown of TgATG9-mAID, bradyzoites were also concurrently treated with 500 µM IAA or equal volume ethanol for 3 days. The CytoID Autophagy Detection Kit 2.0 (Enzo, ENZ-KIT175) was used to stain autolysosomes within live bradyzoites for 45 min prior to fixation with 4% paraformaldehyde, following the manufacturer’s instructions. Fixed coverslips were mounted using ProLong™ Gold Antifade Mountant (Thermo Scientific, P36930). Images were taken on a Zeiss Axio Observer Z1 inverted microscope at 63× and analyzed using Zen 3.7 blue edition software.

### Live cell lattice light sheet microscopy (LLSM)

ME49/TIR1, ME49/TIR1/tdTomato-TgATG8/HaloTag-FYVE, and triple-fluorescent (ME49/TIR1/TgATG9-mAID/tdTomato-TgATG8/HaloTag-FYVE) *T. gondii* strains were differentiated into bradyzoites for 7 days in 35-mm dish, No. 1.5 coverslip, 14-mm diameter, tissue culture dishes (MatTek, P35G-1.5-14-C) in phenol red free alkaline-stress medium consisting of modified RPMI-1640 without phenol red or NaHCO_3_ (Sigma-Aldrich, R8755) supplemented with 3% heat-inactivated fetal bovine serum, 50 mM HEPES, 50 U/mL of penicillin-streptomycin and adjusted to pH 8.2-8.3 with NaOH. After 7 days differentiation, bradyzoites were treated with 500 µM IAA or an equal volume of ethanol as a vehicle control for 24 h for conditional knockdown of TgATG9-mAID. Prior to live cell imaging, bradyzoites were incubated with phenol red free alkaline-stress medium containing 200 nM Janelia Fluor® HaloTag® 646 ligand (Promega, GA1120) for 15 min to label HaloTag-FYVE. After the incubation period, fresh phenol red free alkaline-stress medium was added.

Live imaging of *in vitro T. gondii* cysts was performed using a ZEISS (Carl Zeiss Inc., Oberkochen, Germany) lattice lightsheet 7 microscope (44.83X/1.0 NA Objective at 60° angle to the cover glass, Pco.edge 4.2 CLHS sCMOS camera). The whole cyst volume was imaged in 20 s for a total duration of 5 min with an acquisition of 15 ms exposure for 488 nm, 561 nm, and 640 nm excitations at individual imaging depth. Collected volumes were cropped in Zen 3.5 (Carl Zeiss Inc., Oberkochen, Germany). Images were deconvolved using Zen’s built in deconvolution tool. The deconvolved images were then deskewed in Zen to transform the skewed images into an orthogonal coordinate system based on the coverslip geometry.

### Quantification of lattice light sheet microscopy (LLSM)

Deconvolved and deskewed volumes were imported into Arivis Vision4D 4.0 (Carl Zeiss Inc., Oberkochen, Germany). A linear rotation was applied to each volume and the volumes were further cropped to reduce the data size prior to preprocessing.

Determination of background fluorescence was calculated. Due to inherent parasite autofluorescence and slight bleed through from tdTomato into the 488-nm channel, we used data captured from the non-mNeonGreen-expressing cysts to control for this and to more accurately distinguish true mNeonGreen signal in strains expressing TgATG9-mAID. Control data were preprocessed with Arivis’ Particle Enhancement denoising function to suppress local background and enhance puncta brightness. Enhanced puncta were then segmented with the Blob Finder segmentation tool and the segments were then filtered by volume. Resulting segments were then used to measure the mean intensity of the raw unprocessed control data. Measurements were then exported and mean background values for each time point were calculated. Background volumes were then created in Fiji (v2.9.0/1.53t).

Analysis of experimental data (triple-fluorescent bradyzoites with or without IAA treatment) was subdivided into four Arivis pipelines: In pipeline 1, background volumes created in Fiji were imported and subtracted from the 488-nm channel of the experimental data. The Particle Enhancement function was applied to each channel and the Blob Finder tool was used to segment the resulting volumes. In pipeline 2, cysts were segmented using Arivis’ Machine Learning Segmenter, which utilized a model that was trained on the experimental data. In pipeline 3, puncta segments generated with the Blob Finder function were filtered by volume to eliminate segments smaller than 0.2 um^3^. The remaining puncta segments were then further filtered by their overlap (greater than 90%) as a proxy for co-occurrence of puncta segments. In pipeline 4, filtered puncta segments were then grouped based on overlap with segments from each channel. Object volumes were also calculated for puncta segments from each channel.

Per cyst, the ratio of co-occurrence was calculated from dividing the number of puncta with overlap by the total number of puncta in each channel. This accounts for the overall decreased number of puncta in the 488-nm channel observed in the IAA-treated condition due to TgATG9-mAID knockdown. The co-occurrence between TgATG9 and TgATG8 was determined by taking the number of TgATG9 puncta found to be overlapping with TgATG8 and dividing by the total number of TgATG9 present. The co-occurrence between TgATG8 and TgATG9 was determined by taking the number of TgATG8 puncta found to be overlapping with TgATG9 and dividing by the total number of TgATG8 present. The co-occurrence between TgATG9 and FYVE was determined by taking the number of TgATG9 puncta found to be overlapping with FYVE and dividing by the total number of TgATG9 present. The co-occurrence between TgATG8 and FYVE was determined by taking the number of TgATG8 puncta found to be overlapping with FYVE and dividing by the total number of TgATG8 present.

### Bradyzoite viability assay

The viability of bradyzoites was assessed by combining plaque assay and qPCR normalization of parasite genome numbers [15,25]. *T. gondii* tachyzoites were differentiated into bradyzoites in 6-well tissue culture plates for 7 days with daily media changes. Following differentiation, bradyzoites were treated with 500 µM IAA or an equal volume of ethanol as a control with daily media and treatment changes for 7 days. Following the treatment period, bradyzoites were harvested by pepsin treatment (0.026% pepsin in 170 mM NaCl and 60 mM HCl, final concentration), and filtration [58]. Bradyzoites were enumerated and 500 or 2000 bradyzoites per well were added in triplicate to fresh 6-well tissue culture plates containing confluent HFFs in D10 medium. Bradyzoite-derived plaques were allowed to form undisturbed for 14 days, grown at 37°C under 5% CO_2_. Plaques were counted using a light microscope and plates were then stained with crystal violet fix solution (0.2% of crystal violet and 70% of EtOH) for 15 min at room temperature, rinsed with water. Next, 500 µL of the initial pepsin-treated bradyzoites was used for genomic DNA extraction with the DNeasy Blood and Tissue Kit (Qiagen, 69506). qPCR was performed using 2 µL of genomic DNA in SsoAdvancedTM Universal SYBR® Green Supermix (Bio-Rad, 172-5271) and primers for a 529-base pair repetitive element of *T. gondii* (forward AGGAGAGATATCAGGACTGTAG; reverse GCGTCGTCTCGTCTAGATCG) [78]. The qPCR reactions were performed with the CFX96 Touch Real-Time PCR Detection System (Bio-Rad) using the following parameters: 3 min at 98°C, and 40 cycles of 15 s at 98°C, 30 s at 58.5°C, and 30 s at 72°C. A standard curve was built with 6.4, 32, 160, 800, 4000, 2000 parasite genomes. The number of plaques was normalized to the calculated number of genomes present in the inoculating samples.

### Statistical analyses

Data were analyzed using GraphPad prism. For each data set, outliers were identified and removed using ROUT with a *Q* value of 0.1%. Data were then tested for normality (D’Agostino-Pearson omnibus normality test) and equal variance. Student’s *t*-test or one-way analysis of variance (ANOVA) was performed for normally distributed data with equal or assumed equal variance, when appropriate. Power analysis was not performed when the study was designed. Sample sizes were determined based on the capacity of each assay to collect sufficient values needed to support rigorous analyses and statistical tests.

## Abbreviations

AID: auxin-inducible degron
CFUs: colony-forming units
IAA: indole-3-acetic acid
LLSM: lattice light sheet microscopy
mAID: minimal auxin-inducible degron
PLVAC: plant-like vacuolar compartment.

## Acknowledgements

We thank the Thorner lab at the University of California, Berkeley for providing the Pgk1 antibody, the Sibley lab at Washington University in St. Louis for the TgActin antibody, and the Bradley lab at UCLA for the ATRX1 antibody. We thank the Weiss lab at Albert Einstein College of Medicine for the HaloTag plasmid. We thank Dr. Christophe-Sebastien Arnold for helping in structural modeling of TgATG9. We thank Carruthers lab members: Dr. My-Hang Huynh, Dr. Einar Olafsson, Patrick Rimple, and Tracey Schultz for their key technical support. We thank Biorender for providing a user-friendly tool for creating schematic illustrations.

## Funding

This work was supported by grants from the US National Institute of Health, including R01 AI120607 (V.B.C.), R21 AI160610 (V.B.C.), T32 AI007528 (P.T.), T32 AI007863 (P.T.), F30AI169762 (P.T.), and GM131919 (D.J.K.).

## Disclosures

We declare no competing interests.

## Notes

### Competing Interest Statement

The authors have declared no competing interest.

