## Supplemental Materials for "TgATG9 is required for autophagosome biogenesis and maintenance of chronic infection in *Toxoplasma gondii*"

**Supplemental Material**

**Methods for supplemental figures**

***Microscopy of ScAtg9 and TgATG9 localization in yeast***

Yeast cells were grown to OD600 ~ 0.5 in YPD media and shifted to nitrogen starvation medium for autophagy induction. Images were captured on a Leica DMi8 microscope with a 100X objective and CMOS microscope camera. Eight Z-sections with 0.4-μm spacing between neighboring sections were taken for each view. The image stacks were deconvolved, projected, and manually analyzed by LAS X software.

***TgATG9 phosphorylation mutant generation***

Phosphorylation sites for TgATG9 were determined by searching phosphoproteomics data on the *Toxoplasma* database (ToxoDB) [48]. A total of six residues were annotated to be phosphorylated (S165, S933, S1316, S1332, S1377, S1482). A plasmid was generated containing the last two exons of TgATG9 with S933A, S1316A, S1332A, S1377A, S1482A mutations and a C-terminal 3X-HA tag. A guide RNA targeting upstream of S933 was generated using primers P11/P12 and the guide RNA used for TgATG9-mAID tagging was used. A homology-directed repair template was generated using oligos P13/P14 for PCR amplification of the plasmid containing mutated sites. ME49 ectopically expressing tdTomato-TgATG8 [15], were co-transfected with 100 µg of each guide RNA and 50 µg repair template. At 48 h after transfection, positively transfected parasites were selected using 5 µg/mL phleomycin prior to isolating clones by limiting dilution. Individual parasite clones were validated by PCR amplification using P15/P16 and sent for sequencing with P17/P18/P19. After single-clone isolation, the C-terminal phosphorylation mutant (S933A, S1316A, S1332A, S1377A, S1482A) was used to generate the complete TgATG9 phosphorylation mutant (S165A). A guide RNA targeting upstream of S165 was generated using primers P20/P12 and a guide RNA targeting downstream of S165 was generated using primers P21/P12. A homology-directed repair template was generated by annealing oligos P22/P23. Parasites were co-transfected with 100 µg of each guide RNA and 50 µg repair template. At 48 h after transfection, positively transfected parasites were selected using 5 µg/mL phleomycin prior to isolating clones by limiting dilution. Individual parasite clones were validated by PCR amplification using P24/P25 and sent for sequencing with P26.

***TgATG9 colocalization immunofluorescence assays and quantification***

Starting with *T. gondii* Pru/TgATG9-3XHA parasites [25], Pru/tdTomato-TgATG8/TgATG9-3XHA parasites were generated through transfection with 100 µg of plasmids containing pTubulin-tdTomato-TgATG8-CAT [20] linearized with PmeI for single crossover integration into the tubulin locus. Transfected parasites were selected with chloramphenicol (Sigma-Aldrich, C0378) starting 24 h post-transfection. Single clones were isolated by limiting dilution. Individual parasite clones (Pru/tdTomato-TgATG8/TgATG9-3XHA) were validated by imaging for tdTomato fluorescence.

*T. gondii* Pru/TgATG9-3XHA or Pru/tdTomato-TgATG8/TgATG9-3XHA tachyzoites were differentiated into bradyzoites in 6-well tissue culture plates with 22 mm x 22 mm No. 1.5 coverslips (Globe Scientific, 1404-15) for 7 days. Bradyzoites were fixed with 4% paraformaldehyde, permeabilized with 0.1% Triton X-100 in PBS for 10 min and blocked with 10% FBS in PBS (with 0.01% Triton X-100) for 30 min. Primary and secondary antibodies, along with other staining reagents, were diluted in wash buffer (1% FBS, 1% normal goat serum [Gibco, 16210072], and 0.01% Triton X-100 in PBS). All antibody incubations were performed at room temperature for 1 h. Primary antibodies used were rat anti-HA (Roche, 11867423001; 1:2500), rabbit anti-TgCPL (Carruthers lab; 1:500), and mouse anti-ATRX1 (Bradley lab, UCLA; 1:2000). Secondary antibodies used were goat anti-rat Alexa Fluor 488 (Invitrogen, A-11006; 1:1000), goat anti-rabbit Alexa Fluor 594 (Invitrogen, A-11012; 1:1000), and goat anti-mouse Alexa Fluor 594 (Invitrogen, A-11032; 1:1000). Coverslips were mounted using ProLong™ Gold Antifade Mountant (Thermo Scientific, P36930). Images were taken on a Zeiss Axio Observer Z1 inverted microscope at 63× and analyzed using Zen 3.7 blue edition software.

Quantification of colocalization was conducted using CellProfiler 4.2.5 software. Individual cysts were manually outlined based on phase-contrast images and saved as “Objects”. Colocalization within identified Objects was calculated using the “MeasureColocalization” function with a threshold setting for the top 25% of the intensity in each channel. The Manders (M1) colocalization coefficients were calculated, exported, and plotted as shown in Figure S3B. At least 30 cysts were analyzed per biological replicate (n = 3).

**Supplemental Figure 1.**

**Figure S1.** ScAtg9 and TgATG9 proteins are expressed in yeast and colocalize with RFP-Atg8. Fluorescence microscopy images containing a widefield view of yeast cells expressing RFP-Atg8 (*atg1∆ atg9∆* RFP-Atg8) and ScAtg9-mNG or TgATG9-mNG. Cells were imaged after 1 h (top panels) and 3 h (bottom panels) nitrogen starvation. Arrows indicate examples of colocalization between mNG and RFP-Atg8. *atg1∆* cells were used as this mutant results in Atg9 restriction to the phagophore assembly site. While TgATG9-mNG is expressed in these cells, the qualitative appearance of colocalization between TgATG9 and RFP-Atg8 is different from ScAtg9 and RFP-Atg8 congruent with the differences in their respective abilities to rescue bulk autophagy (Figure 2) likely due to defects in efficient cycling of TgATG9.

**Supplemental Figure 2.**

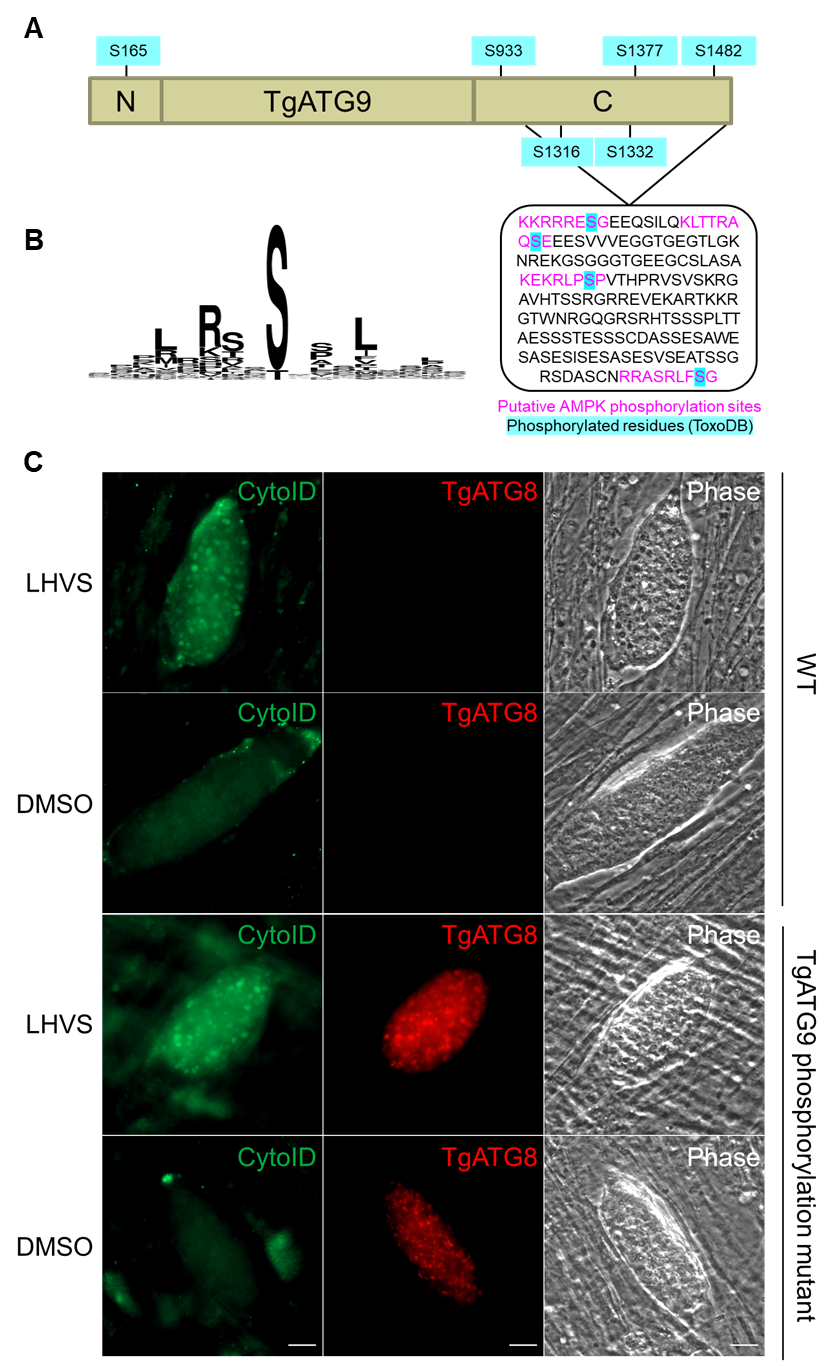

**Figure S2.** TgATG9 phosphorylation does not regulate its function in autophagy. (**A**) TgATG9 is annotated in the Toxoplasma informatics database (ToxoDB) to be phosphorylated at 6 sites (serines, S) with 4 sites on the C terminus containing putative AMPK recognition motifs (magenta). (**B**) Logo motif of 64 known AMPK phosphorylation sites in humans. The logo motif was generated using WebLogo. (**C**) Autolysosome staining (CytoID) of in vitro WT (ME49) or TgATG9 phosphorylation mutant (ME49/tdTomato-TgATG8/TgATG9 S165A, S933A, S1316A, S1332A, S1377A, S1482A) differentiated cysts treated with LHVS or DMSO for 72 h. A WT control that did not express tdTomato-TgATG8 was used to confirm tdTomato expression in the TgATG9 phosphorylation mutant. Scale bars: 10 μm.

**Supplemental Figure 3.**

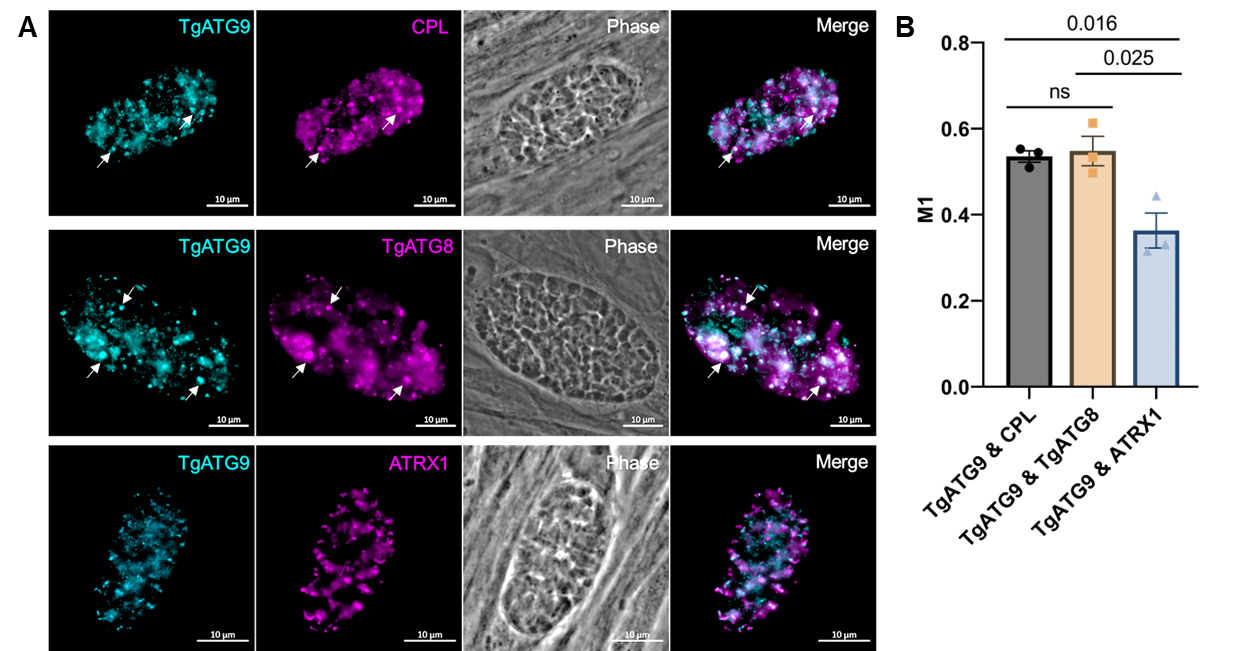

**Figure S3.** TgATG9 colocalizes with the parasite’s digestive organelle (PLVAC) and autophagosomes in bradyzoites. (**A**) Immunofluorescence staining of in vitro differentiated cysts for colocalization of TgATG9-3xHA and the PLVAC marker CPL (top panel), autophagosome marker TgATG8 (middle panel), and apicoplast marker ATRX1 (bottom panel). Scale bars: 10 mm. (**B**) Quantification of M1 colocalization coefficients using CellProfiler software. Bars represent mean ± SEM. Statistical analysis was done using ordinary one-way ANOVA with Tukey’s multiple comparison tests, p-values denoted above bars (n=3 biological replicates).

**Supplemental Video 1.**

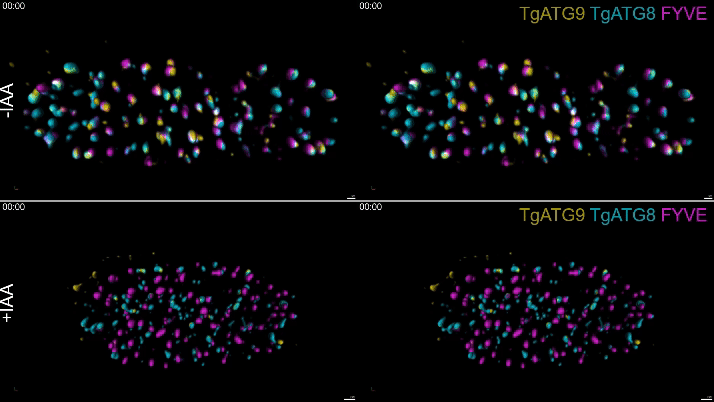

**Video S1.** Videos of representative in vitro differentiated cysts imaged with LLSM treated with IAA or vehicle control (-IAA) for 24 h. Scale bars: 2 μm (top row) and 5 μm (bottom row). Each cyst is shown over a 5-min time course in both single planar (left) and rotational views (right). Volumetric renderings for TgATG9, TgATG8, and TgFYVE were generated in Arivis Vision4D software. Videos are from the same cysts shown in Figure 4B.

**Supplemental Video 2.**

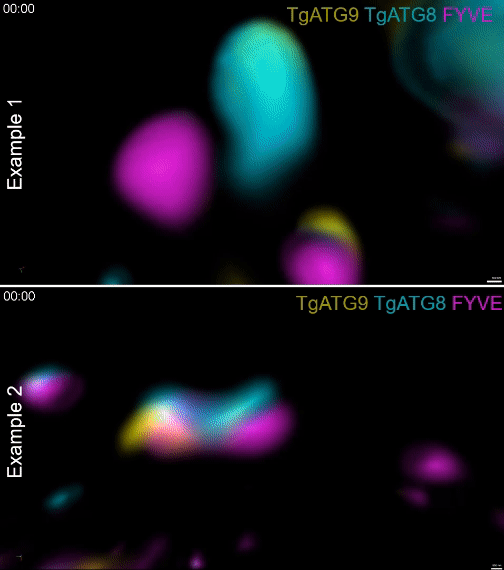

**Video S2.** Videos of representative examples of dynamic associations between TgATG9, TgATG8, and TgFYVE imaged with LLSM in untreated parasites (-IAA). Videos are from the same examples shown in Figure 4C.
